# Quantitative genetics and modularity in cranial and mandibular morphology of *Calomys expulsus*

**DOI:** 10.1101/008151

**Authors:** Guilherme Garcia, Rui Cerqueira, Erika Hingst-Zaher, Gabriel Marroig

## Abstract

Patterns of genetic covariance between characters (represented by the covariance matrix ***G***) play an important role in morphological evolution, since they interact with the evolutionary forces acting over populations. They are also expected to influence the patterns expressed in their phenotypic counterparts (***P***), because of limits imposed by multiple developmental and functional restrictions on the genotype/phenotype map. We have investigated genetic covariances in the skull and mandible of the vesper mouse (*Calomys expulsus*) in order to estimate the degree of similarity between genetic and phenotypic covariances and its potential roots on developmental and functional factors shaping those integration patterns. We use a classic *ad hoc* analysis of morphological integration based on current state of art of developmental/functional factors during mammalian ontogeny and also applied a novel methodology that makes use of simulated evolutionary responses. We have obtained ***P*** and ***G*** that are strongly similar, for both skull and mandible; their similarity is achieved through the spatial and temporal organization of developmental and functional interactions, which are consistently recognized as hypothesis of trait associations in both matrices.

## Introduction

Morphological integration refers to the interconnection among morphological elements due to genetic, functional and developmental relationships between such elements, as expressed during the course of development (Olson and Miller 1958; Cheverud 1996). From a classical quantitative genetics perspective (Falconer and Mackay 1996), these interconnections among morphological elements are represented as phenotypic and genetic covariance or correlation matrices (***P*** and ***G***, respectively). The structure represented by ***G*** is the product of additive effects of multiple *loci*, affecting multiple traits through pleiotropy and linkage disequilibrium. Phenotypic covariance or correlation structure is then defined as the result of the interactions between these genetic effects and environmental perturbations. However, genes act upon phenotypes through development, which can be thought as a function that maps genotypes into phenotypes (Wagner 1984; Polly 2008). In this context, integration refers to the structure of gene-trait mapping. Theoretical considerations made by Wagner (1996) suggest that this genotype/phenotype map would display a modular organization; i.e., that traits are grouped in subsets, and each trait subset is affected by a subset of genes with pleiotropic effects mostly confined to each group.

While ***G*** is associated with the additive portion of genetic variation, which can be understood as the linear approximation of this developmental function centered at the mean phenotype (Wagner 1984), its structural aspects are also dependent upon non-linear developmental factors, traditionally viewed in quantitative genetics as dominance and epistasis (Wolf et al 2001). Developmental dynamics involves not only the interaction among many genes, but the interactions among the developing cells and tissues, both in terms of differential expression and signaling, and mechanical interactions; these complex interactions often lead to non-linear effects within the genotype/phenotype map (Turing 1952; Polly 2008; Krupinski et al 2011; Tiedemann et al 2012; Watson et al 2013). In the mammalian skull, for instance, there is a series of overlapping steps that define the adult phenotype, such as neural crest cell migration, the formation of cell condensations of osteoblasts and the subsequent differentiation of these cells into osteocytes, brain growth and muscle-bone interactions (Hallgrímsson and Lieberman 2008; Franz-Odendaal 2011); therefore, while development has a tendency to produce covariance among morphological elements, the pattern described by phenotypic or genetic covariance matrices is often difficult to categorize, due to the superposition of these effects in shaping covariance patterns (Hallgrímsson et al 2009; Roseman et al 2009). Furthermore, developmental dynamics may limit the full expression of genetic variation (Hallgrímsson et al 2009), since mutational effects may destabilize the system as a whole; hence, developmental dynamics may act as internal stabilizing selection (Cheverud 1984) and, in equilibrium, the genetic covariance structure will match those pattern arising both from stabilizing selection, drift, and mutational effects (Lande 1980).

Empirical evidence on the association between morphological traits and their underlying genetic architecture focused on the identification of pleiotropic quantitative trait *loci* supports the hypothesis of modular organization in gene-trait associations. For instance, the partitioning of the skull into Face and Neurocranium (Leamy et al 1999) and partitioning of the mandible into Alveolar Process and Ascending Ramus (Cheverud et al 1997; Mezey et al 2000; Cheverud 2006); however, Klingenberg et al (2004) has found evidence contrary to these mandibular partitions, although the disagreement between these results might arise due to methodological differences between traditional and geometric morphometrics. Without a strict adherence to the view of a modular genotype/phenotype map, Roseman et al (2009) found a positive association between correlation among pairs of traits and their amount of shared pleiotropic effects.

The structural aspects expressed in ***G*** are central to all discussions regarding the evolution of complex phenotypes since ***G*** mediates the response to directional selection (Lande 1979), imposing several properties of such response in a microevolutionary scale (Hansen and Houle 2008), with possible consequences on a macroevolutionary scale (Marroig and Cheverud 2005). For example, the cosine of the angle between a given selection gradient (***β***; the vector of fitness slopes over phenotypes; Lande 1979; Falconer and Mackay 1996) and the response it produces (***Δz̄***), called flexibility (*f*) by Marroig et al (2009), is affected both by the imposed selection direction and the covariance structure of the trait system considered. For example, in a system composed of three traits (Figure 1), if directional selection is aligned with the genetic covariance structure in a population (which describes morphological integration), response to such selection will be direct (Figure 1a-b), yielding a high flexibility value for this selection gradient. However, if the selection gradient is unaligned to population covariance structure, response to selection will be deflected from the optimal direction (Figure 1c-d), lowering the associated flexibility value.

**Fig. 1.**
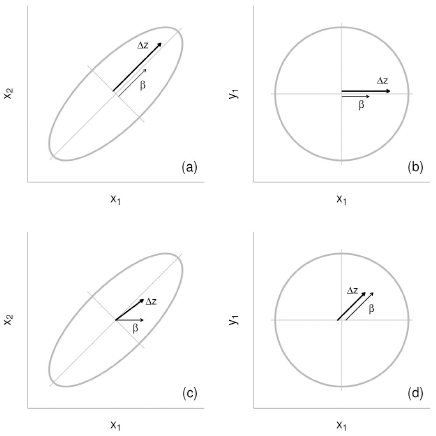
Theoretical expectations regarding the relationship between selection gradients and response to selection with respect to covariance structure. In this system composed of three traits, the covariance between *x*_1_ and *x*_2_ is positive, while both these traits are independent from *y*_1_. In (a) and (b), the ***β*** applied over the covariance matrix between *x*_1_, *x*_2_ and *y*_1_ matches the covariance matrix structure; therefore, ***Δz̄*** follows ***β*** closely, yielding a high *f* value. In (c) and (d), ***β*** does not match the structure embedded in the covariance matrix; hence, ***Δz̄*** is deflected due to the covariance between *x*_1_ and *x*_2_ (c); in this case, the associated *f* value is lower.

The relationship between selection gradients and response to selection may also be affected by the line of genetics least resistance (LLR; Schluter 1996), the first eigenvector of ***G***, which represent the major axis of multivariate genetic variation on a population for a given trait system. If genetic variation is concentrated in this direction in a given population (Figure 2a), response to selection will be biased towards it; only selection vectors nearly orthogonal to the LLR would escape such bias. Hence, the population would exhibit overall lower flexibility values, compared to another population with less variation concentrated along the LLR (Figure 2b). With respect to mammalian morphological systems, the LLR is, in most cases, associated with size variation (Marroig et al 2004; Porto et al 2009, 2013), and can be thought as a global integrating factor, since it impacts morphological variation as a whole. Therefore, the effect of this source of variation over morphological integration has to be taken into account when dealing with such systems (Marroig et al 2004; Mitteroecker and Bookstein 2007; Porto et al 2013).

**Fig. 2.**
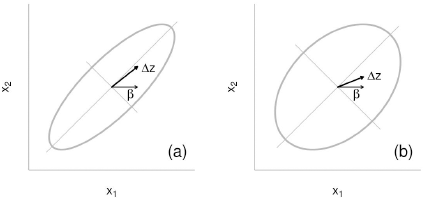
Expectations regarding the relationship between flexibility values and variation along the LLR. In (a) and (b), the ***β*** applied over the covariance matrix between *x*_1_ and *x*_2_ only selects for average increase in *x*_1_. In both situations, response to selection is deflected due to the covariance between the two traits. However, the response is more strongly biased in (a) than in (b), leading to a lower associated *f* value for the first situation.

Due to the importance of the genetic covariance structure in multivariate trait systems, there has been a consistent effort in estimating ***G*** (Steppan et al 2002). However, such matrices are often difficult to estimate, since this estimation depends on the availability of genealogical relationship information within a population (Falconer and Mackay 1996; Lynch and Walsh 1998). Furthermore, even if such information exists, the estimation of ***G*** is severely prone to error (Hill and Thompson 1978; Meyer and Kirkpatrick 2008; Marroig et al 2012), since their estimation is usually based on sample units (families) that are an order of magnitude lower than sample sizes for ***P*** (Roff 1997).

A solution to this limitation was proposed by Cheverud (1988), who compared a wide range of ***P***s and ***G***s estimated mostly from morphological traits, finding similarities in covariance patterns, especially for morphological traits. Therefore, the so called ‘Cheverud's Conjecture’ (Roff 1995) states that the patterns expressed by ***G*** will be mirrored by their phenotypic counterparts, given that the covariance between environmental effects (***E***) are either low, uncorrelated or similar to ***G*** in their structure. While it may seem odd that environmental and genetic covariance patterns would have similar structures, both sources of variation exert their effects over phenotypic covariance structure through common developmental pathways (Klingenberg 2008). Several authors (e.g.: Roff 1995, 1997; Reusch and Blanckenhorn 1998; Roff and Fairbairn 2011; Dochtermann 2011) have formally tested this conjecture for different traits systems across a wide range of taxa, finding supporting evidence on its favor.

In the present article, we estimated phenotypic and genetic covariance and correlation matrices for both cranial and mandibular traits for a population of the vesper mouse *Calomys expulsus*, a sigmodontine rodent, comparing these matrices with respect to the correspondence of patterns predicted by Cheverud's Conjecture (Cheverud 1988; Roff 1995). We also investigated the association between covariance/correlation structure and hypotheses of trait associations based upon functional and developmental relationships, using the traditional methods established by several authors (Cheverud 1995; Marroig and Cheverud 2001; Marroig et al 2004; Porto et al 2009) and a novel methodology, based on the expectations regarding the relationship between selection gradients that represent hypotheses of trait association and response to selection to such gradients, as suggested by Hansen and Houle (2008).

## Materials and Methods

### Sample

The genus Calomys (Muroidea, Cricetidae) consists of six species of small sigmodontine rodents that occurs in open and forested areas across Central South America (Hershkovitz 1962; Bonvicino and Almeida 2000; Bonvicino et al 2003). Molecular phylogenetic analysis indicates that *Calomys* is a basal clade of the Phylottini tribe (Steppan et al 2004). The vesper mouse, *Calomys expulsus*, occurs over the dry biomes of Central Brazil in sympatry with the delicate vesper mouse (*C. tener*) at the southern limit of its distribution. Recently, both morphological, karyotypic and molecular data confirm the validity of the species (Bonvicino and Almeida 2000; Almeida et al 2007).

In order to estimate covariance matrices for cranial and mandibular traits in *C. expulsus*, we used a series of specimens deposited at the mammal collection of the Museu Nacional do Rio de Janeiro (MNRJ). This series constitute a captive-bred colony originated from 37 sexually immature individuals captured in a single locality in the State of Minas Gerais, Brazil (Fazenda Canoas, 16°50′S, 43°35′W, 800m). Twenty-one males and 16 females were placed under controlled conditions and randomly paired, producing over 400 sibs in a one-year period; each litter was maintained with its dam until weaning. During this period, couples were rearranged at random; therefore, the colony design is unbalanced, containing both paternal and maternal half-sibs. Our sample is constituted of 365 skulls and 228 mandibles of individuals from this colony whose genealogical information is available, comprising individuals of both sexes and of different age classes.

### Landmarks and measurements

We registered three-dimensional coordinates for 20 cranial landmarks (Figure 3) using a Microscribe MX digitizer (Microscribe, IL). We registered bilaterally symmetrical landmarks in both sides, when available, for a total of 32 landmarks. Details of the digitizing procedure and landmark definition can be found in Cheverud (1995). We calculated a set of 35 inter-landmark distances (Figure 3); trait names follow the landmarks of which they are composed. We measured each individual twice, in order to access measurement error through the estimation of repeatability (Lessels and Boag 1987). After this estimation, we averaged repeated measures, thus reducing our measurement error (Falconer and Mackay 1996); we also averaged distances that are present on both sides of the skull. Therefore, our set of 35 cranial traits is comprised of these averaged inter-landmark distances. Our inter-landmarks distances are designed to measure individual bones, thus capturing localized aspects of the skull development and avoiding the shortcomings of dealing with full length measures capturing several bones at same time (skull length for example) or principal components of shape.

**Fig. 3.**
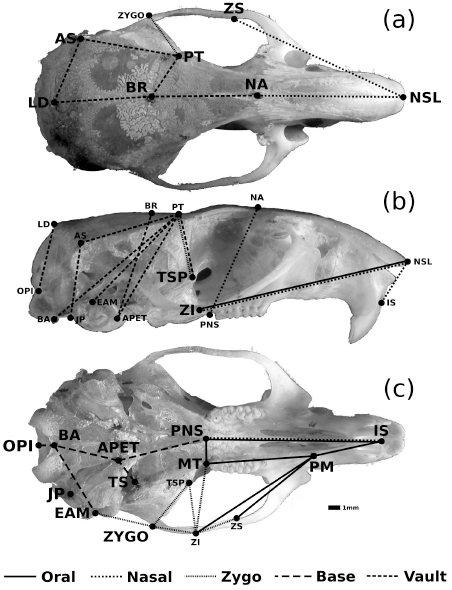
The 20 registered landmarks represented over an adult skull of *C. expulsus* in dorsal (a), lateral (b) and ventral (c) views. Landmark names are highlighted in the particular view at which they are best defined. The set of 35 inter-landmark distances are also represented; different line types represent distinct modularity hypotheses to which traits are associated. Some distances (ISPNS, NSLZI, PTTSP) are associated with more than one hypothesis, indicated by double lines.

We registered bi-dimensional coordinates for 10 mandibular landmarks (Figure 4) over pictures we took of both hemimandibles (when available) using tpsDig2 (Rohlf 2006), and calculated a set of 20 inter-landmark distances (Figure 4). We averaged measures from both hemimandibles to compose our set of 20 mandibular traits; as with the skull, trait names follow the landmarks of which they are composed. Since the mandible is a single bone, both landmark placements and inter-landmark distances suffer from a intrinsic uncertainty, because most landmarks are defined as the limits of mandibular processes (Bookstein 1991); therefore, our mandibular traits may not be as biologically meaningful as our cranial traits, since they only describe mandibular morphology, not being associated with particular developmental processes.

**Fig. 4.**
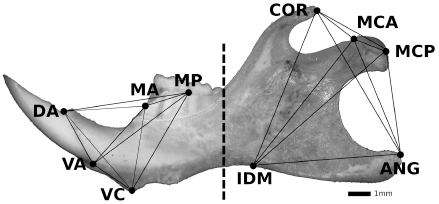
The 10 registered landmarks represented over an adult left hemimandible of *C. expulsus*. Lines between landmarks represent the set of 20 Euclidean distances calculated between landmarks; the dashed line represents the two distinct morphological integration hypotheses tested over these traits, the (anterior) Alveolar Processes and the (posterior) Ascending Ramus. Landmark definitions follow: **DA**: dorsal incisor alveolus; **MA**: anterior limit of m1; **MP**: posterior limit of m1; **COR**: coronoid process dorsal limit; **MCA**: anterior limit of mandibular condyle; **MCP**: posterior limit of mandibular condyle; **ANG**: angular process posterior limit; **IDM**: maximum dorsal inflection between angular and alveolar processes; **VC**: ventral limit of chin; **VA**: ventral incisor alveolus.

We took a second set of pictures of a subset of 30 individuals; we used this subset to estimate repeatability taking into account both measurement error and the error associated with picture registration. To this purpose, we collected four sets of data for these individuals, two from each set of pictures. Repeatability in this case is the percentage of variance explained by the individual factor alone, excluding variation from both measurement and photo acquisition errors.

### Estimation of covariance matrices

Since sexual dimorphism and ontogenetic variation are of little interest within the present context, we explored such sources of variation with respect to our sets of 35 cranial traits and 20 mandibular traits through analysis of multivariate covariance (MANCOVA), and evaluated the significance of these factors (and their interaction term) using Wilks’ *λ* test. In order to remove the impact these effects may have on covariance structure, we used the residual covariance matrices of the two separate linear models (one for cranial traits and a second one for mandibular traits) as estimates of phenotypic covariance matrices for the *C. expulsus* population. We also estimated ***P***s for different ages (20, 30, 50, 100, 200, 300 and 400 days) and for males and females. These matrices were compared in order to test whether these matrices are similar and can be grouped into a single ***P*** for the entire sample.

### **G**-matrix estimation

For the estimation of genetic covariance matrices of these two sets of traits, we use the approach proposed by Runcie and Mukherjee (2013) that uses a Bayesian mixed-effects model in order to produce ***G*** estimates. This model builds upon the simple animal model (Lynch and Walsh 1998) by defining both ***G*** and ***P*** as depending on a set of latent traits, i.e., linear combination of the original set of traits. The prior distributions that define these factors impose two important constraints on their structure: first, that there is a declining influence of subsequent factors in terms of amount of explained phenotypic and genetic variance; second, the prior distributions impose sparsity over each factor; therefore, each latent trait is associated to a limited number of traits in the original set. Therefore, based on this factorization, ***G*** can be represented as

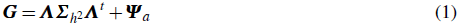

where ***Λ*** is the matrix of trait loadings on each factor, 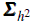 is the diagonal matrix of factor heritabilities, and ***Ψ***_*a*_ is the diagonal matrix of trait-specific genetic variances. Due to its characteristic decomposition of ***G***-matrices, this model was named Bayesian Sparse Factor analysis of Genetic covariances (BSFG) by Runcie and Mukherjee (2013).

In order to obtain posterior distribution samples for ***Λ***, 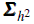, and ***Ψ***_*a*_ in both cranial and mandibular trait sets, we used a MCMC algorithm, obtaining 1000 samples from 10000 iterations, with a burnin period of 1000 iterations. We initialized the MCMC run with 17 latent traits for both cranial and mandibular trait sets. We did a handful of MCMC runs with different initial prior distribution parameters, but posterior distributions were not affected by these differences. For each sample taken from the posterior distribution for those parameters described above, we constructed a different ***G***, obtaining a posterior distribution for this matrix; we also estimate a posterior mean matrix, which is our best estimate for ***G***.

Using the posterior distribution of heritabilities associated with each factor (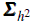), we test whether any given factor has a genetic variance different from zero, by computing the highest posterior density (HPD) intervals for all diagonal elements of 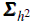. Using the posterior sample of ***G***-matrices, we also estimated posterior intervals for trait heritabilities.

We also estimated ***G*** using a classical REML algorithm (Shaw 1987), treating genetic variances for each trait and pairwise genetic covariances as independent estimates; we grouped these estimates in matrix form to compose a ***G*** estimate. We did such proceeding using both a REML algorithm we wrote using R ({R Core Team} 2013) and also in WOMBAT (Meyer 2007), estimating additive variances and covariances using the simple animal model (Lynch and Walsh 1998). We compared these estimates with those obtained from the Bayesian sparse factor analysis outlined above; however, we chose to use the Bayesian model for four reasons. First, it produces estimates for the entire ***G*** structure simultaneously; when using REML algorithms, this structure has to be divided into its components (genetic variances and covariances) to be computationally tractable. Second, since ***G*** estimates from the BSFG model are samples from the posterior distribution, they are constrained to have only positive eigenvalues, even if only a reduced number of latent traits is included in the model. Furthermore, the latent traits (***Λ***) obtained from the BSFG model might be informative in terms of the underlying processes influencing the genotype/phenotype map (*sensu* Wagner and Altenberg 1996). Finally, the posterior distribution for ***G*** we obtained from the model can be used to obtain posterior intervals for all matrix parameters we estimated (see the following subsections for details).

Although we prefer the BSFG model estimates for ***G***, the conventional REML estimates can be used to estimate the effective sample size (*N*_*eff*_) of each *h*^2^ estimate; this represents the approximate number of independent additive values used to estimate that particular *h*^2^ value (Cheverud 1995). The relationship

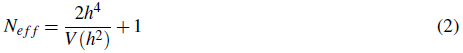

can be used to estimate effective sample sizes, where *h*^4^ is the squared *h*^2^ estimate, and *V* (*h*^2^) is the estimated variance of the estimate, considering a normal approximation to the likelihood profile for that estimate. The geometric mean of individual *N*_*eff*_ estimates for each trait can then be used as a proxy for the experiment-wise effective sample size for ***G*** as a whole (Cheverud 1995).

*Size Variation* In order to investigate the influence of size variation over morphological integration, we also obtained matrices whose variation associated with the first principal component has been removed, as this component is usually associated with size in mammalian morphological systems (Wagner 1984; Marroig et al 2004; Porto et al 2013). As suggested by Bookstein et al (1985), a residual matrix whose size variation has been removed can be obtained using the following relationship:

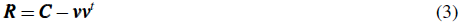

where ***C*** represents the raw (size retained) matrix and ***v*** denotes the unstandardized first eigenvector of ***C***; its norm is equivalent to the square root of the associated eigenvalue.

### Matrix Comparisons

We compared our estimated ***P*** with the mean ***G*** estimated from the BSFG model, by using the Random Skewers method (Cheverud and Marroig 2007) for covariance matrices and matrix correlation followed by Mantel's test for significance (Cheverud et al 1989) for correlation matrices. We also compared ***P*** with all ***G*s** from the posterior distribution of the BSFG model in order to estimate uncertainty in matrix similarity, obtaining a distribution of average Random Skewers and matrix correlation values. These comparisons directly tests Cheverud's Conjecture (Cheverud 1988; Roff 1995) of similarity between genetic and phenotypic covariance/correlation patterns.

We also compared covariance matrices using the Selection Response Decomposition method (Marroig et al 2011), which pinpoints differences between matrices with respect to trait covariance structure. Using this method, we are able to observe the correspondences between phenotypic and genetic covariance patterns for both mandibular and cranial traits. We compared both matrices with size variation retained and removed using the SRD method.

### Morphological Integration Analysis

In order to investigate whether patterns described by ***P*** and ***G*** in our sets of cranial and mandibular traits conform to hypothesis of trait association due to functional and developmental interactions, we measured both magnitude and pattern of integration in phenotypic and genetic covariance/correlation matrices. We represented magnitude of morphological integration using the ICV (Shirai and Marroig 2010), which is the coefficient of variation of eigenvalues in a given covariance matrix.

We took two different approaches to investigate patterns of integration. In the classical approach (Olson and Miller 1958; Cheverud 1995), we constructed theoretical matrices to represent hypotheses of association between traits due to functional and/or developmental relationships expected *a priori* (Cheverud 1995). The hypotheses for cranial traits followed those proposed by Porto et al (2009) for the Order Rodentia (Figure 3). We constructed a total of nine theoretical matrices for cranial traits: five associated with localized hypotheses (Oral, Nasal, Zygomatic, Vault and Base), two associated with more global hypotheses that contrast early and late developmental patterns in mammals (Neurocranium and Face, respectively), and two composite hypotheses associated with these two groups (Total and Neuroface, respectively). Notice that Total integration correspond to the sum of all five individual hypothesis into one composite hypothesis and the Neuroface corresponds to the conjoint test of late and early developmental influence (or the sum of both individual hypothesis, Face and Neurocranium). With respect to mandibular traits, we considered the distinction between the Alveolar Processes and the Ascending Ramus, according to Figure 4, as proposed by several authors (e.g.: Cheverud et al 1997; Klingenberg et al 2004; Willmore et al 2009). We also tested a third, composite hypothesis (Total), corresponding to the sum of Alveolar and Ascending hypotheses. We calculated matrix correlations using Pearson-product moment correlation between each constructed theoretical matrix and the correlation matrices estimated from data in order to estimate the association between them (Cheverud et al 1989; Cheverud 1995). We estimated significance for each correlation calculated this way by Mantel's test. Notice that, in the context of testing for hypotheses of association between traits, this classical approach is equivalent to a simple test of differences between the averages of two groups, integrated *vs.* non-integrated traits, as a Student's t-test, albeit with the significance test modified (using Mantel) to account for the non-independence of the observations (in this case, correlations among traits).

We investigated pattern and magnitude of morphological integration in matrices with size variation retained and removed; in order to compare estimates between these two types of matrices, we used the modularity index proposed by Porto et al (2013), calculated as the difference between the mean correlation taken from the set of correlations bounded by a given hypothesis and the mean correlation taken from the complementary set of correlations, divided by the ICV calculated for the corresponding covariance matrix. This index is, therefore, comparable across matrices with different magnitudes of integration, which is the case when comparing matrices with and without size variation.

By using the ***G*** posterior distribution obtained from the BSFG model, we estimated posterior intervals for both ICV and modularity indexes for each hypothesis. By estimating these intervals, we intend to represent the error associated with these parameters estimated over our mean ***G***. It also allows us to compare the estimated parameter values for both ***P*** and ***G***, under the null hypothesis that differences between these parameters for the two types of matrices are only due to error in estimating the mean ***G***. This comparison addresses the possibility of local differences between ***P*** and ***G***, in a manner similar to the SRD method.

#### Simulated Evolutionary Responses

The second and novel approach used to test hypotheses of trait associations is based on simulated evolutionary responses to selection, as suggested *en passant* by Hansen and Houle (2008). We used all covariance matrices available, including those whose size variation has been removed (***P*** and ***G*** for both skull and mandible). For each matrix, we obtained an empirical distribution of flexibility (Marroig et al 2009) without any *a priori* assumptions. Using 10,000 random normalized vectors drawn from a multivariate normal distribution without correlation structure, we estimated mean value (*f̄*) and the 95% confidence interval for *f*.

Using this *f* distribution, we explored our hypotheses of trait associations (as suggested by Hansen and Houle 2008) by constructing theoretical selection gradients that represent our hypothetical modules. Each ***β*** associated with a given hypothesis has a value of one for traits within the hypothetical module and zero otherwise; afterwards, each vector is also normalized. For each ***β*** constructed in this fashion (Oral, Nasal, Zygomatic, Vault, Base, Face and Neurocranium for the skull; Alveolar and Ascending for the mandible), we estimated the associated *f* values; the Neuroface (for the skull) and Total (for both skull and mandible) hypotheses cannot be properly represented as selection gradients, and are therefore excluded from this analysis. In order to test whether each of these vectors corresponds to a set of integrated traits in a given covariance matrix, we compare these *f* values with the critical 95% interval around *f̄* for that matrix, obtained from selection gradients that are random with respect to that matrix covariance structure. If that particular *f* is higher than the critical value from the distribution, we considered this as evidence that the associated ***β*** represents trait associations embedded in that covariance matrix. In this case, the corresponding ***Δz̄*** follows ***β*** more closely than expected by chance alone; therefore, covariance structure interferes to a lesser extent in the response to selection in this case, demonstrating that the population has, to some degree, independent variation in the direction of that particular ***β***.

In a manner similar to the ICV and modularity index, the ***G*** posterior distribution allows us to estimate posterior distributions for both *f̄* and *f*-values associated with hypotheses of morphological integration. These distribution intend to represent the error associated with estimation of flexibilities in ***G***-matrices; hence, they allow us to compare *f*-values between ***P*** and ***G***, under the null hypotheses that differences in these values are only due to error in estimating ***G***.

#### BSFG Factors and Morphological Integration

In order to investigate the relationship between the factors estimated by the BSFG model and our hypotheses of morphological integration, we calculated vector correlations between these factors and the vectors constructed to represent our hypotheses, as outlined in the previous section. We estimated significance for these correlations by using a distribution of correlations obtained from random vectors, in a manner similar to the estimation of significance in the Random Skewers method (Cheverud and Marroig 2007). If vector correlations between any given combination of latent traits and hypothesis vectors is higher than the critical value estimated from random vector correlations, we consider that correlation as evidence of non-random association between latent traits and hypothesis.

## Results

### Measurement Error Assessment

Repeatabilities for cranial traits (Table S1) ranged from 0.654 to 0.998 with an average value of 0.956 and standard deviation 0.068. For mandibular traits (Table S2), values ranged from 0.869 to 0.992, with mean 0.961 and s.d. 0.037. Traits with low repeatabilities were those with low variances, such as MTPNS. For cranial traits, these results for repeatability should not impact other results, since repeatabilities for averaged traits that have been measured twice (as we have done here) are higher than traits with single measurements (Falconer and Mackay 1996). For mandibular traits, the lowest repeatability value is above 0.85; therefore, we expect that further analysis should not be impacted by the error arising from both photo acquisition and measurement procedure.

### Matrix Estimation

Regarding estimation of phenotypic matrices, both linear models adjusted for removal of age and sex fixed effects (and their interaction) were significant for all factors used, for both sets of traits; separate estimates for ***P*** at different ages and in both males and females are also similar (all comparisons above 0.68; results not shown). Therefore, we used the residual covariance matrices from these models as estimates of ***P***.

Heritabilities for latent traits recovered from the BSFG model (Table S3) indicate that, for both cranial and mandibular trait sets, four of the 17 latent traits have posterior distributions of *h*^2^ that do not include the null value. For both cranial and mandibular trait sets, the first latent trait recovered is a direction associated with size variation, as indicated by the negative sign associated with almost all traits in ***λ***_1_ for cranial traits (Table 1, with the exception of BRPT and BAOPI) and the positive sign associated with ***λ***_1_ for mandibular traits (Table 2). Considering the posterior distribution of factor loadings in ***λ***_1_ for both skull and mandible, almost all traits have factor loadings different from zero, except for the cranial trait BRPT and mandibular traits VCVA and MCAMCP.

**Table 1.** Unstandardized first principal components obtained from covariance matrices and factors retrieved from the BSFG model for cranial traits. Bold values indicate those factor loadings that differ from zero, according to the 95% posterior interval of factor loadings.

| Trait | Hypothesis | Region | $p_1$ | $g_1$ | $\lambda_1$ | $\lambda_2$ | $\lambda_3$ | $\lambda_4$ |
| --- | --- | --- | --- | --- | --- | --- | --- | --- |
| ISPM | Oral | Face | -0.868 | -0.268 | **-0.347** | 0.013 | -0.003 | 0.003 |
| PMZS | Oral | Face | -1.196 | -0.37 | **-0.464** | **0.132** | 0.012 | -0.009 |
| PMZI | Oral | Face | -1.47 | -0.469 | **-0.582** | **0.167** | 0.031 | -0.001 |
| PMMT | Oral | Face | -0.764 | -0.23 | **-0.302** | 0.001 | -0.008 | -0.053 |
| MTPNS | Oral | Face | -0.085 | -0.023 | **-0.031** | 0.002 | 0.011 | -0.003 |
| ISPNS | Oral/Nasal | Face | -1.804 | -0.551 | **-0.708** | 0.095 | 0.005 | **-0.14** |
| NSLZI | Oral/Nasal | Face | -2.607 | -0.815 | **-1.029** | **0.19** | 0.01 | 0.009 |
| ISNSL | Nasal | Face | -0.563 | -0.181 | **-0.235** | 0.017 | -0.005 | 0 |
| NSLNA | Nasal | Face | -1.822 | -0.55 | **-0.699** | 0.09 | -0.044 | -0.021 |
| NSLZS | Nasal | Face | -2.22 | -0.68 | **-0.858** | **0.194** | -0.001 | 0.006 |
| NAPNS | Nasal | Face | -0.879 | -0.269 | **-0.356** | -0.012 | -0.006 | **-0.095** |
| PTZYGO | Zygomatic | Face | -0.951 | -0.296 | **-0.4** | **-0.202** | 0 | 0.016 |
| ZSZI | Zygomatic | Face | -0.454 | -0.154 | **-0.2** | -0.027 | 0.008 | 0.002 |
| ZIMT | Zygomatic | Face | -0.854 | -0.274 | **-0.349** | **0.102** | -0.011 | 0.023 |
| ZIZYGO | Zygomatic | Face | -0.474 | -0.137 | **-0.2** | 0.005 | 0.003 | -0.009 |
| ZITSP | Zygomatic | Face | -0.851 | -0.262 | **-0.342** | 0.03 | -0.022 | -0.001 |
| EAMZYGO | Zygomatic | Face | -0.504 | -0.144 | **-0.191** | 0.001 | -0.021 | 0.005 |
| ZYGOTSP | Zygomatic/Vault | Face/Neuro | -0.68 | -0.221 | **-0.287** | 0.019 | 0.004 | 0.007 |
| PTTSP | Vault | Neurocranium | -0.382 | -0.116 | **-0.155** | -0.036 | 0.003 | -0.011 |
| NABR | Vault | Neurocranium | -0.797 | -0.25 | **-0.321** | **0.146** | -0.058 | 0.017 |
| BRPT | Vault | Neurocranium | 0.001 | 0.016 | 0.014 | **-0.06** | -0.017 | 0.002 |
| BRAPET | Vault | Neurocranium | -0.556 | -0.16 | **-0.219** | **-0.109** | **0.047** | -0.032 |
| PTAPET | Vault | Neurocranium | -0.906 | -0.272 | **-0.371** | **-0.177** | -0.002 | 0.009 |
| PTBA | Vault | Neurocranium | -1.414 | -0.43 | **-0.572** | -0.16 | 0.016 | 0.042 |
| PTEAM | Vault | Neurocranium | -1.13 | -0.344 | **-0.468** | **-0.243** | 0.002 | 0.025 |
| LDAS | Vault | Neurocranium | -0.148 | -0.04 | **-0.052** | -0.001 | **0.081** | 0.002 |
| BRLD | Vault | Neurocranium | -0.437 | -0.122 | **-0.152** | -0.017 | **0.369** | -0.001 |
| OPILD | Vault | Neurocranium | -0.69 | -0.212 | **-0.284** | **-0.094** | **-0.065** | -0.014 |
| PTAS | Vault | Neurocranium | -0.741 | -0.22 | **-0.291** | -0.173 | **0.085** | 0.016 |
| JPAS | Vault | Neurocranium | -0.627 | -0.192 | **-0.262** | **-0.107** | -0.006 | 0.004 |
| PNSAPET | Base | Neurocranium | -1.015 | -0.319 | **-0.415** | **0.069** | 0 | **0.169** |
| APETBA | Base | Neurocranium | -0.688 | -0.211 | **-0.271** | 0.048 | 0.002 | 0.013 |
| APETTS | Base | Neurocranium | -0.168 | -0.052 | **-0.069** | -0.006 | -0.001 | -0.019 |
| BAEAM | Base | Neurocranium | -0.496 | -0.155 | **-0.2** | **0.039** | 0.016 | 0.008 |
| BAOPI | Base | Neurocranium | 0.139 | 0.048 | **0.06** | -0.011 | 0.003 | -0.012 |

**Table 2.** Unstandardized first principal components obtained from covariance matrices and factors retrieved from the BSFG model for mandibular traits. Bold values indicate those factor loadings that differ from zero, according to the 95% posterior interval of factor loadings.

| Trait | Hypothesis | $p_1$ | $g_1$ | $\lambda_1$ | $\lambda_2$ | $\lambda_3$ | $\lambda_4$ |
| --- | --- | --- | --- | --- | --- | --- | --- |
| DAMA | Alveolar | -0.127 | -0.078 | **0.170** | 0.006 | 0.015 | **0.053** |
| DAMP | Alveolar | -0.162 | -0.096 | **0.196** | 0.007 | 0.016 | -0.047 |
| MAMP | Alveolar | -0.036 | -0.018 | **0.024** | 0 | 0 | **-0.105** |
| DAVC | Alveolar | -0.142 | -0.067 | **0.115** | 0.003 | **0.128** | **-0.058** |
| MAVC | Alveolar | -0.174 | -0.085 | **0.156** | 0.003 | 0.034 | 0.040 |
| MPVC | Alveolar | -0.194 | -0.103 | **0.186** | -0.007 | -0.031 | -0.034 |
| DAVA | Alveolar | -0.076 | -0.043 | **0.080** | 0.003 | **0.141** | -0.026 |
| MAVA | Alveolar | -0.141 | -0.080 | **0.162** | 0.006 | -0.001 | **0.072** |
| MPVA | Alveolar | -0.168 | -0.094 | **0.178** | 0 | **-0.04** | -0.044 |
| VCVA | Alveolar | -0.063 | -0.021 | 0.029 | 0.001 | -0.009 | **-0.036** |
| CORMCA | Ascending | -0.160 | -0.090 | **0.070** | **-0.263** | -0.002 | -0.019 |
| CORMCP | Ascending | -0.212 | -0.108 | **0.093** | **-0.284** | 0.001 | 0.004 |
| MCAMCP | Ascending | -0.050 | -0.013 | 0.019 | -0.001 | 0.004 | 0.028 |
| CORANG | Ascending | -0.387 | -0.180 | **0.238** | **-0.167** | -0.011 | -0.026 |
| MCAANG | Ascending | -0.234 | -0.109 | **0.168** | -0.013 | -0.003 | -0.021 |
| MCPANG | Ascending | -0.183 | -0.083 | **0.120** | 0.004 | -0.009 | -0.027 |
| CORIDM | Ascending | -0.364 | -0.152 | **0.257** | 0.033 | 0.011 | -0.021 |
| MCAIDM | Ascending | -0.360 | -0.169 | **0.258** | -0.068 | 0.013 | -0.036 |
| MCPIDM | Ascending | -0.390 | -0.169 | **0.250** | -0.058 | 0.013 | -0.017 |
| ANGIDM | Ascending | -0.413 | -0.182 | **0.278** | -0.062 | 0.010 | -0.020 |

The second cranial latent trait (***λ***_2_ in Table 1) has positive loadings associated with most facial traits, and negative loadings related to neurocranial traits. Therefore, this factor can be understood as a contrast between these two regions. The third factor, ***λ***_3_, is only associated with Vault characters, while the fourth, ***λ***_4_, recovers a pattern similar to ***λ***_2_, involving contrasts between facial and neurocranial traits located in the ventral side of the skull.

The three remaining mandibular latent traits (***λ***_2_, ***λ***_3_, ***λ***_4_ in Table 2) have only a handful of traits whose factor loadings are different than zero, and such sparse factor loadings have localized effects with respect to mandibular partitioning. The second factor affect only traits belonging to the Ascending Ramus, while the other two factors are associated with Alveolar traits.

Following Equation 1, we constructed our estimated mean ***G*** and the posterior distribution for ***G*** using all factors we estimate. Comparing the mean ***G*** estimated by the BSFG model with ***G***s estimated using REML (both our own R-based-algorithm and Wombat) using Random Skewers (for covariance matrices) and matrix correlation followed by Mantel's test (for correlation matrices) indicate that matrices estimated from the three methods are fairly similar. Random Skewers correlations ranged from 0.84 to 0.91, and matrix correlations ranged from 0.72 to 0.84; all correlations are indicative of a lack of structural dissimilarities under their respective tests of significance (for ***P***(*α*) *<* 10^−4^). Estimated trait heritabilities are also similar among these three different types of estimation methods (Figures S1 and S2). Therefore, the BSFG estimates of ***G*** we use here, for both cranial and mandibular trait sets are very similar to the traditional REML estimates.

Using the REML estimates for *h*^2^, we were able to estimate the effective sample sizes for both cranial (Table S1) and mandibular (Table S2) traits. For cranial traits, we obtained an average *N*_*eff*_ of 20 individuals; for mandibular traits, the estimated average *N*_*eff*_ is 32 individuals.

For both cranial and mandibular matrices, the first eigenvector of both ***P*** and ***G*** are indeed size vectors (represented as ***P***_1_ and ***G***_1_ in Tables 1 and 2). Hence, by removing the first eigenvector of these matrices following Equation 3, we obtain matrices without size variation, removing both variation from scaling and allometry.

### Matrix Comparisons

All comparisons between ***P*** and the mean ***G***, using both Random Skewers correlations for covariance matrices and matrix correlation followed by Mantel's test, reject the null hypothesis of structural dissimilarity between these matrices (Table 3), with very high correlation values (all above 0.9). Comparing ***P*** with any ***G*** derived from the posterior distribution of the BSFG model results in matrix correlations which reject this null hypothesis for all comparisons. Correlation values for covariance matrices are systematically higher than those derived from correlation matrices, considering both the comparison between ***P*** with the mean ***G*** and the posterior distribution of matrix correlations (Table 3).

**Table 3.** Comparison among ***P*** and ***G*** for both cranial and mandibular traits. “Value” represents the average Random Skewers correlation or matrix correlation *Γ*-values between ***P*** and the mean ***G*** for covariance and correlation matrices, respectively. Other statistics (Mean, Median, 95% Posterior Interval) refer to the distribution of the same matrix correlation statistics between ***P*** and the posterior distribution for ***G***. All comparisons reject the null hypothesis of no structural similarity between compared matrices at ***P***(*α*) = 0.001.

| | | Value | Mean | Median | $PI-$ | $PI+$ |
| --- | --- | --- | --- | --- | --- | --- |
| Skull | Cov | 0.988 | 0.955 | 0.963 | 0.902 | 0.987 |
|  | Cor | 0.975 | 0.927 | 0.936 | 0.856 | 0.969 |
| Mandible | Cov | 0.960 | 0.919 | 0.923 | 0.869 | 0.963 |
|  | Cor | 0.942 | 0.832 | 0.839 | 0.731 | 0.920 |

The comparison between ***P*** and ***G*** using the SRD method (Figure 5) indicates that, in matrices whose size variation has been retained, few traits have marked differences in covariance structure. In the cranial set (Figure 5a; with an average SRD score of 0.99), only trait BRPT shows a substantial difference in trait covariance structure between ***P*** and ***G***; in the mandibular set (Figure 5c; with an average SRD score of 0.98), trait MCAMCP has the lowest SRD score. Comparing matrices whose size variation has been removed (Figures 5b and 5d) yields lower SRD scores (although still quite high: 0.92 for the skull, and 0.89 for the mandible); differences between the two matrices are more distributed through each set.

**Fig. 5.**
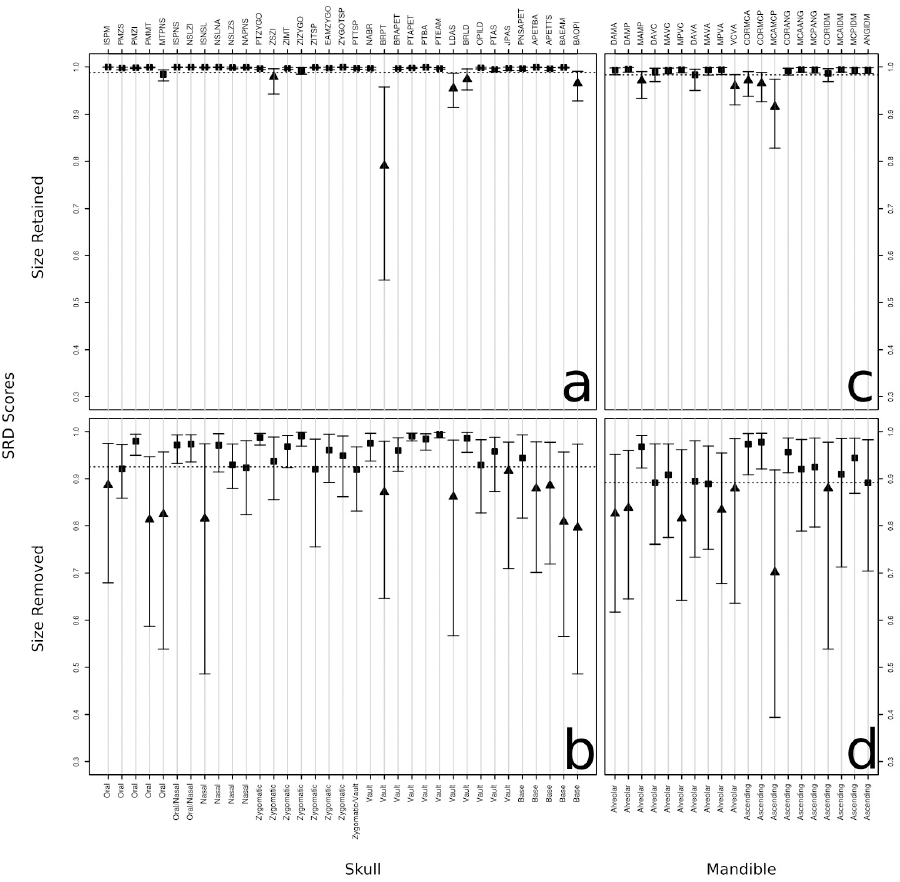
Selection Response Decomposition plots for the comparison between phenotypic and genetic matrices for both cranial (a-b) and mandibular (c-d) traits; size variation has either been retained (a, c) and removed (b, d) for each matrix in these comparisons. In all comparisons, dashed lines represent average SRD scores. Cranial or mandibular traits indicated with triangles are those which differ significantly between each ***P***/*G* set, with ***P***(*α*) = 0.025.

### Morphological Integration

When considering matrices whose size variation has been retained, ICV's estimated for both cranial and mandibular trait sets do not differ between ***P*** and ***G***, as both phenotypic ICV's are within the 95% posterior interval constructed using sampled ***G***s (Table 4). For size-free matrices, for cranial traits, ICV values between ***P*** and ***G*** also do not differ; however, the phenotypic value for the mandibular ***P*** ICV (1.332) is below the lower bound of the posterior interval (1.576). It is also noteworthy that, when size is removed, there is a substantial reduction in ICV values in both trait sets.

**Table 4.** Magnitude and Pattern of phenotypic and genetic integration, as measured by ICV and Modularity Index, respectively. Both cranial and mandibular matrices are represented, with size variation either retained or removed. Posterior Intervals (*PIγ*=0.95) are associated with the posterior distribution for ***G***, representing parameter uncertainty in ***G*** parameters. Bold values are associated with morphological integration hypotheses that are recognized in a given matrix by Mantel's test (*P*(*α*) = 0.05); italic values are marginally significant (*P*(*α*) = 0.1).

|  | Size | Retained |  |  |  | Removed |  |  |  |
| --- | --- | --- | --- | --- | --- | --- | --- | --- | --- |
| | | $\mathbf{P}$ | $\mathbf{G}$ | $PI-$ | $PI+$ | $\mathbf{P}$ | $\mathbf{G}$ | $PI-$ | $PI+$ |
| Skull | ICV | 4.091 | 3.794 | 2.615 | 4.791 | 1.502 | 1.448 | 1.299 | 2.042 |
|  | Oral | *0.049* | *0.056* | 0.031 | 0.087 | **0.018** | **0.025** | 0.008 | 0.037 |
|  | Nasal | **0.083** | **0.092** | 0.065 | 0.127 | 0.016 | *0.024* | 0.007 | 0.037 |
|  | Zygomatic | -0.005 | -0.005 | -0.031 | 0.025 | -0.013 | -0.015 | -0.026 | 0.004 |
|  | Vault | -0.011 | -0.009 | -0.024 | 0.017 | **0.035** | **0.045** | 0.015 | 0.067 |
|  | Base | -0.058 | -0.056 | -0.073 | -0.039 | -0.005 | 0.001 | -0.016 | 0.018 |
|  | Total | 0.007 | 0.011 | 0.002 | 0.029 | **0.021** | **0.029** | 0.010 | 0.046 |
|  | Face | **0.039** | **0.045** | 0.032 | 0.059 | 0.002 | 0.004 | -0.001 | 0.009 |
|  | Neuro | -0.031 | -0.032 | -0.043 | -0.02 | **0.017** | **0.019** | 0.006 | 0.031 |
|  | Neuroface | **0.010** | **0.014** | 0.007 | 0.025 | **0.014** | **0.017** | 0.005 | 0.027 |
| Mandible | ICV | 2.737 | 2.656 | 2.283 | 3.120 | 1.332 | 1.822 | 1.576 | 2.355 |
|  | Alveolar | 0.003 | 0.011 | -0.019 | 0.049 | **0.055** | **0.027** | 0.002 | 0.054 |
|  | Ascending | *0.050* | 0.033 | -0.010 | 0.078 | **0.056** | **0.040** | 0.009 | 0.060 |
|  | Total | **0.039** | **0.031** | 0.006 | 0.060 | **0.080** | **0.048** | 0.011 | 0.079 |

Regarding pattern of morphological integration in cranial traits (Table 4), both Nasal and Facial trait subsets are identified by Mantel's test; the composite Neuroface hypothesis is also identified. When size variation is removed, the Oral, Vault and Neurocranial regions are identified; the Neuroface region is identified again. There is agreement in highlighted regions between ***P*** and ***G***; modularity indexes estimated for phenotypic matrices are within the range of their respective 95% posterior intervals, indicating that patterns of morphological integration are not different between ***P*** and ***G***, when considering the uncertainty associated with estimating ***G***.

Considering the mandible, only the Total composite hypothesis is identified by Mantel's test in matrices with size variation (Table 4). When removing size variation, all three hypotheses (Alveolar, Ascending and Total) are identified. As with the cranial trait set, there is agreement between the hypothesis identified for both phenotypic and genetic matrices. The posterior distribution of modularity indexes indicate, for those matrices with size variation retained, that patterns of morphological integration are not different between matrices compared; however, when considering the comparison between matrices without size variation, modularity indexes estimated for ***P*** for both Alveolar (0.055) and Total (0.08) hypotheses are slightly above the upper limits of their respective posterior intervals (0.054 and 0.079).

#### Flexibility

Regarding our analysis of morphological integration based on simulated evolutionary responses (Table 5), for both cranial matrices, the ***β***s associated with Facial, Oral, and Nasal traits had flexibility values significantly higher than expected by chance alone; when size variation is removed, the ***β***s associated with the Vault and Base were also higher than average *f* values. For mandibular traits, the ***β*** associated with the Ascending Ramus had *f* values higher than *f̄* for both ***P*** and ***G***; when size variation is removed, the Alveolar Process ***β*** was also associated with a high *f* value in ***P***, but not in ***G***.

**Table 5.**
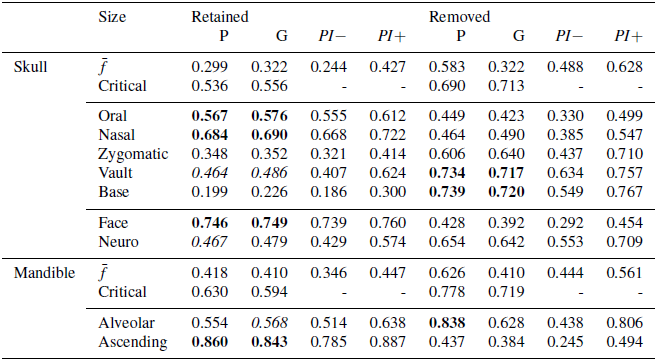
Flexibilities of phenotypic and genetic matrices, Both cranial and mandibular matrices are represented, with size variation either retained and removed. Posterior Intervals (*PIγ*=0.95) are associated with the posterior distribution for ***G***, representing parameter uncertainty in ***G*** parameters. Bold values are associated with morphological integration hypotheses that are recognized in a given matrix by comparison with the null distribution of flexibilities for any given matrix (*P*(*α*) = 0.05); italic values are marginally significant (*P*(*α*) = 0.1). ‘Critical’ refers to the 95% upper bound value for these comparisons.

Considering the differences in flexibility values between ***P*** and ***G***, only the *f*-value estimated for the Alveolar region in ***P*** with size removed (0.838) is above the upper bound of the posterior distribution of *f*-values derived from sampled ***G***s (0.806); the *f̄* value calculated for ***P*** (0.626) is also above the limit drawn by the posterior distribution (0.561). Remaining *f*-values, in all other phenotypic matrices, are within the range of their respective posterior intervals.

#### Factors and Morphological Integration

Correlations between factors recovered from the BSFG model and our hypotheses (Table 6) indicate that the first factor for cranial traits has high correlation values with the Oral, Nasal and Facial ***β***s, while the second factor is mildly correlated with the Vault ***β***. Regarding mandibular latent traits, ***λ***_1_ has high correlations with both Alveolar and Ascending *β*s, while ***λ***_2_ has a high correlation with only the ***β*** associated with the Ascending Ramus.

**Table 6.**
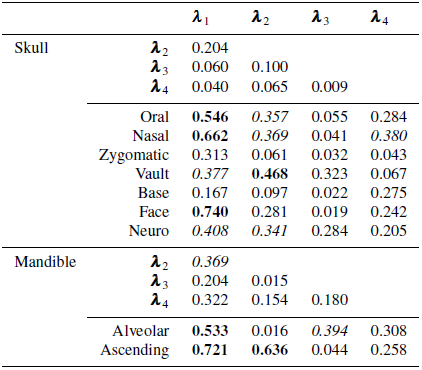
Correlation between factors recovered from the BSFG model and morphological integration hypotheses. Values highlighted represent vector correlation values higher than those expected by chance, with ***P***(*α*) = 0.05 for italic values and ***P***(*α*) = 0.01 for bold values.

| | | $\lambda_1$ | $\lambda_2$ | $\lambda_3$ | $\lambda_4$ |
| --- | --- | --- | --- | --- | --- |
| Skull | $\lambda_2$ | 0.204 | | | |
| | $\lambda_3$ | 0.060 | 0.100 | | |
| | $\lambda_4$ | 0.040 | 0.065 | 0.009 | |
|  | Oral | **0.546** | *0.357* | 0.055 | 0.284 |
|  | Nasal | **0.662** | *0.369* | 0.041 | *0.380* |
|  | Zygomatic | 0.313 | 0.061 | 0.032 | 0.043 |
|  | Vault | *0.377* | **0.468** | 0.323 | 0.067 |
|  | Base | 0.167 | 0.097 | 0.022 | 0.275 |
|  | Face | **0.740** | 0.281 | 0.019 | 0.242 |
|  | Neuro | *0.408* | *0.341* | 0.284 | 0.205 |
| Mandible | $\lambda_2$ | *0.369* | | | |
| | $\lambda_3$ | 0.204 | 0.015 | | |
| | $\lambda_4$ | 0.322 | 0.154 | 0.180 | |
|  | Alveolar | **0.533** | 0.016 | *0.394* | 0.308 |
|  | Ascending | **0.721** | **0.636** | 0.044 | 0.258 |

## Discussion

The results we obtained here point out that phenotypic and genetic covariance structure are very similar in the vesper mouse, for both skull and mandible; while differences in trait-specific covariances exist, as shown by the SRD method (Figure 5), these differences are local and do not affect the overall covariance structure, as shown by matrix correlation between ***P*** and ***G*** (Table 3), and also by average SRD scores, even when the effects of size variation is controlled (Figure 5). Hence, these comparisons corroborate Cheverud's Conjecture (Cheverud 1988; Roff 1995) of similarity between patterns of correlation/covariance structure in both ***P*** and ***G***. We will now focus on exploring the potential underlying causes of this similarity.

The observed differences in trait-specific covariances, pinpointed by the SRD method (Figure 5) do not affect our hypotheses of trait associations attributed to developmental and functional interactions. Overall, the same subsets of traits are integrated in both phenotypic and genetic correlation (Table 4) and covariance (Table 5) patterns. Although these results agree between each ***P***/***G*** set, there are minor differences between the traditional method employed on correlation matrices and the method we propose here, used on covariance matrices. For instance, while the Neurocranium is identified as a valid partition in correlation matrices without size variation, the same partition is not identified by our evolutionary simulations approach.

While these differences might occur due to differences in type II error rates for both tests (which can be subject to future investigations), our simulation-based approach has the advantage of being directly related to the use of covariance matrices in evolutionary quantitative genetics theory. The classical test of morphological integration (Cheverud et al 1989) has its roots on the comparison of distance matrices (Mantel 1967), while our simulation approach is based on the analysis of evolutionary response to selection as suggested by Hansen and Houle (2008). Therefore, this approach is more appropriate for dealing with systems (and representations of such systems, i.e., covariance matrices) that could be subjected to evolutionary change. On the other hand, the classical approach has the advantage of being able to handle more complex hierarchical structures that cannot be properly represented as a vector. For example, the contrast between early and late developmental factors, represented by a composite hypothesis that compares correlations within facial and neurocranial traits with correlations between these two groups, i.e. the Neuroface hypothesis, cannot be tested in the selection gradient framework, since this test considers the correlation structure of all traits simultaneously and cannot be represented as a binary vector. Therefore, both approaches complement each other, being under different premises and expectations.

The distribution of posterior ***G*** statistics that represent pattern and magnitude of morphological integration (ICV, modularity index, flexibility values) indicate that almost all differences in these values between ***P*** and ***G*** may be attributed only to error in estimating ***G***, with the exception of features associated with the mandibular matrices when size is removed. Considering together results in Tables 4 and 5 indicate that the mandibular ***P*** is less integrated and more modular than its genetic counterpart, as indicated by the lower ICV value and higher *f*-values and modularity indexes associated with Alveolar traits. However, these differences are found only when size variation is removed; such source of variance represents a substantial portion of overall magnitude of integration. Therefore, these differences between ***P*** and ***G*** in the mandible are minor compared to the overall similarity in matrix structure.

### Size Variation and Integration Patterns

Removing size from both phenotypic and genetic matrices produces substantial changes in covariance structure. First and foremost, differences between trait-specific covariances increase (Figure 5). Magnitude of integration, as measured by ICV, drops considerably, in all matrices whose size variation has been removed. Patterns of integration also change; different hypotheses of trait association are recognized as valid.

The variation in size within populations has a pervasive effect throughout development, from zygote to adult. However, growth will have an heterogeneous effect over traits, producing allometric variation. Overall, regarding the mammalian skull, facial traits are more influenced by growth than neural traits during the pre-weaning period and afterwards (Zelditch and Carmichael 1989; Hallgrímsson et al 2009), due to overall growth and muscle-bone interactions associated with both breastfeeding and mastication. Thus, the influence of size variation is more pervasive in the integration of facial traits in the skull (Tables 4 and 5).

Size variation also has an influence on integration patterns of the mandible; the Ascending Ramus is recognized as a valid hypothesis of trait association by our evolutionary simulation approach when we consider matrices whose size variation has been retained (Table 5). The masseter muscle, which affects the growth of the associated Ascending Ramus through its function, has a marked influence on pre-weaning development (Zelditch and Carmichael 1989). The interactions between the masseter and the mandible also produce an extensive reworking of the condylar cartilage (Herring 2011), which might explain the difference in MCAMCP (condyle length) covariance structure between ***P*** and ***G*** (Figure 5). Therefore, both functional and developmental interactions shape the observed integration patterning through growth in both skull and mandible.

When size variation is removed, different features of the developmental system are observed. In the skull, we observe integration in neurocranial traits, in both analysis (Tables 4 and 5). For the mandible, the traditional analysis (Table 4) recognizes both Alveolar and Ascending Ramus hypothesis as valid. These other aspects of integration reflect aspects of prenatal development. In the skull, they are associated with prenatal brain growth (Hallgrímsson and Lieberman 2008; Hallgrímsson et al 2009); in the mandible, the contrast between the two regions may reflect the distinct cell condensation from which each structure arises (Atchley and Hall 1991; Ramaesh and Bard 2003). While trait-specific covariance structure diverges between ***P*** and ***G*** when size is removed (Figures 5b and 5d), these differences do not impact the relationships between these traits, as the same hypotheses of trait associations are recognized in both matrices (Tables 4 and 5). Therefore, these differences in trait-specific covariance structure depicted by the SRD method may reflect differences in covariance structure outside of those regions that are recognized as integrated when size is removed.

Integration patterns produced by size variation and developmental patterning are temporally organized, since their main influence occur at postnatal and prenatal development, respectively. Hence, these developmental factors affect covariance patterns in the skull and mandible with different strengths, with size integration partially masking the effect of other patterns produced by prenatal growth (Hallgrímsson and Lieberman 2008; Hallgrímsson et al 2009). The relative contribution from both these effects to response to selection will reflect this hierarchy, with size variation contributing to a major extent relative to prenatal patterning, acting as a line of least genetic resistance, with evolutionary consequences (Schluter 1996; Marroig and Cheverud 2005).

Our treatment of size variation contrasts with the approach advocated by landmark-based geometric morphometrics (Bookstein 1991; Zelditch et al 2004), in which size is treated as a separate variable, centroid size. Certainly, this approach has advantages; for instance, it allows for a clear representation of allometric relationships, by regressing shape variables on centroid size. With our approach, we are not able to differentiate between isometric and allometric variation; our size factors are associated with both variation components. However, quantitative genetics analyses using landmark-based geometric morphometrics (e.g. Klingenberg and Leamy 2001; Klingenberg et al 2004; Martínez-Abadías et al 2011) have not fully appreciated this advantage, since size variation is treated as a nuisance with respect to shape and its effects are removed or simply ignored. Therefore, the effects of size variation over shape genetic covariance structure remain unexplored under the purview of geometric morphometrics.

However, the main issue regarding the use of landmark-based data to investigate covariance structure lies at the assumption of isotropic variation around each landmark necessary to perform the Generalized Procrustes Superimposition (GPS; Dryden and Mardia 1998; Linde and Houle 2009). Under a morphological integration perspective, we expect that local function and/or developmental factors will produce differences in both magnitude and direction of landmark variation; therefore, the assumption of isotropic variation is incompatible with analyses of covariance structure. If a particular dataset breaks this assumption and GPS is performed nonetheless, the result is that landmark variation will be forced to conform to the assumption by distributing total variation across all landmarks (Linde and Houle 2009) and consequently erasing most of the actual integration patterns.

This issue might explain the contrast between the results from Cheverud et al (1997) and Klingenberg et al (2004) regarding the organization of mandibular pleiotropic QTL effects in the same *Mus* population using traditional and geometric morphometrics, respectively. While Cheverud et al (1997) found that most of these effect are confined within both Alveolar Process and Ascending Ramus (see also Mezey et al 2000; Cheverud 2006), Klingenberg et al (2004) found evidence that most pleiotropic effects are shared by the two regions. Some authors have proposed solutions that circumvent this problem (e.g. Theobald and Wuttke 2006; Linde and Houle 2009; Márquez et al 2012), but these works have not yet been appreciated outside the literature of morphometric theory.

### Latent Traits and Developmental Factors

The latent traits recovered from the BSFG model (Tables 1 and 2) may be understood as a local linear approximation of the developmental function, centered at the mean phenotype, as they are derived from a particular hyper-parametrization of ***G*** we use here (Runcie and Mukherjee 2013). When considering a set of linear approximations to developmental factors, other sets obtained through rotations of the original set may also be considered as a valid approximation (Wagner 1984; Wolf et al 2001; Mitteroecker 2009; Runcie and Mukherjee 2013). However, the set of latent traits we recovered here are constrained by sparsity, and thus are not arbitrary with respect to rotation.

The first factor recovered for both cranial and mandibular traits is associated with size variation (Tables 1 and 2), and reflect the influence of this source of variation over morphological integration. Interestingly, the size factor recovered for both skull and mandible breaks the *a priori* assumption of sparsity made by the BSFG model, which is indicative of strong evidence favoring the existence of such factor. Remaining latent traits reflect our hypotheses of morphological integration (Table 6); in the skull, the second trait recovered is associated with facial and neurocranial traits simultaneously, albeit with opposing signs (Table 1). This factor is usually recovered from mammalian skull variation from our previous studies (e.g.: Marroig et al 2004); in the mandible, each latent trait recovered aside from the first has localized effects over either Alveolar or Ascending traits (Table 2).

The structure of latent traits we recovered here are an abstraction over more complex developmental dynamics; individually, their may or may not reflect actual aspects of the organization of pleiotropic effects. The first factor in both skull and mandible recovers a biologically realistic factor, that is, size variation; in fact, any gene involved in controlling the amount of cell metabolic output or cellular growth and division will contribute to size variation. The second factor in the skull essentially contrasts the face with the neurocranium. While this factor certainly captures one essential aspect of mammalian skull development (one that contrasts early-and late-developmental growth as recognized here in our integration hypotheses) on the other hand it does not correspond to the genetic basis of skull covariation. In fact, most QTLs found in previous studies (Cheverud et al 1997; Leamy et al 1999) affect either the Face (31%) or Neurocranium (31%) separately and only 38% affected the whole skull. More importantly, of those 38% with general effects on the skull only 20% (7.6% out of the total QTLs) had effects with opposite signals on the Face and Neurocranium which would indicate antagonist pleiotropy. For the mandible essentially the same pattern was found with 26% of the 41 QTLs found did affect the whole mandible and the remaining 74% have only localized effects with no evidence of antagonistic pleiotropy. Therefore, latent traits for the skull and mandible display different structures, although there are only minor differences in their underlying genetic architectures with respect to pleiotropic effects, as demonstrated by QTL studies. These latent traits should then be interpreted as a set, and are not to be taken individually as representations of true developmental factors, in the same manner that individual principal components also cannot be regarded as biologically meaningful (Zelditch et al 2004; Adams et al 2011; Berner et al 2011; Berner 2012).

These results suggest that similarity between ***P*** and ***G*** is the result of limits imposed by the developmental system to the action of additive effects over morphological traits (Cheverud 1984; Wolf et al 2001). These limits are the result of both developmental and functional interactions, and are reflected in our hypothesis of trait associations, which are consistently recognized in both matrices and in the latent traits identified by the BSFG model. While these latent traits may be understood as developmental modules (Runcie and Mukherjee 2013), or more likely, as combinations or contrasts between interacting modules, they are temporally organized throughout development (Zelditch and Carmichael 1989; Hallgrímsson and Lieberman 2008), and their effect over both ***P*** and ***G*** will reflect this organization.

It is paramount to understand modularity not as a static feature of morphological systems, but as a feature embedded within a dynamical process, that is, development, which will produce an integrated phenotype (Cheverud 1996; Wagner 1996; Hallgrímsson et al 2009). While our hypothesis of trait associations might be quite simple (as they are essentially binary descriptors of subsets of traits), our results are interpreted in the light of developmental dynamics. Therefore, rather than address only to the identification of modules, one should consider the dynamics of the underlying development when trying to understand both the structure and evolution of complex morphological elements subject to morphological integration.

## Conclusions

Integration and modularity are paramount features of morphological systems, and the evolution of such systems is strongly affected by them. The patterns embedded within genetic and phenotypic covariance matrices capture both these phenomena simultaneously; hence, by estimating covariance patterns within populations, we are able to quantify the relative influence of both magnitude and pattern of integration, and the relationship between ***G*** and ***P***, established by the genotype/phenotype map. Here, we investigate both these levels of organization in the skull and mandible of a *C. expulsus* population, finding remarkable similarities in their overall structure and in the patterns they describe, as predicted by Cheverud's Conjecture (Cheverud 1988; Roff 1995). This similarity between covariance patterns is achieved through the constraints imposed by the developmental system, through the association of morphological elements due to developmental and functional interactions, formulated as hypotheses of trait associations that are recognizable in both genetic and phenotypic covariance structure.

## Acknowledgements

We would like to thank N. P. Barros, A. M. Marcondes, F. Almeida, L. Araripe, J. M. Freschi for help with lab work; F. A. Machado, D. Melo for comments on early drafts. We also thank D. Runcie and S. Murkherjee for help with their BSFG model codes. This work has been supported by grants from CNPq (Conselho Nacional de Pesquisa e Desenvolvimento), FAPERJ (Fundação de Amparo à Pesquisa do Estado do Rio de Janeiro), FAPESP (Fundação de Amparo à Pesquisa do Estado de São Paulo), MMA (Ministério do Meio Ambiente) and MCT (Ministério de Ciência e Tecnologia).

